# Analysis of the possible pathways of cancer cell repopulation after irradiation in vitro

**DOI:** 10.1101/2025.11.12.688027

**Authors:** Y.A. Eidelman, E.O. Shevchenko, S.G. Andreev

**Affiliations:** N.M. Emanuel Institute of Biochemical Physics, Kosygin str. 4, 119334, Moscow, Russia; National Research Nuclear University MEPHI, Kashirskoye shosse 31, 115409, Moscow, Russia

## Abstract

The study of the pathways of cancer stem cell (CSC) repopulation after therapeutic exposure is of great importance for planning effective therapy schemes. In this work, a mathematical model of the proliferation of a population of cancer cells after irradiation is developed, taking into account the presence of three subpopulations (differentiated cancer cells, CSCs, and senescent cells), the plasticity of malignant cells (dedifferentiation, transition of non-stem cells to a stem state), and the transition of cells to a senescent state following irradiation. It is shown that the phenomenon of an increase in the fraction of CSCs after irradiation, observed in the *in vitro* experiment on MCF-7 breast cancer cells, can be explained by symmetrical division of CSCs and dedifferentiation of non-stem cancer cells.

## INTRODUCTION

The use of biological models of cancer cell growth makes it possible to look for ways to increase patient survival by optimizing a large number of physical, biological factors and a variety of clinical data. The development and analysis of models of tumor response to radiation allows to explore different scenarios and concepts underlying tumor resistance for optimal planning of therapeutic intervention schemes [1].

Today, it is generally accepted that cancer recurrence after radiotherapy is caused by the fraction of radiation-resistant cells in tumors. These cells have stem properties, and after therapy or in the course of multi-fraction therapy they repopulate and replace dead tumor cells. Studying the proliferation kinetics of these cells, termed cancer stem cells (CSCs), is of great practical importance. Modern experimental methods make it possible to isolate putative CSCs from the bulk population of cancer cells by the presence of certain markers. For example, in normal and malignant mammary cells, cells with stem properties are characterized by the CD44^+^/CD24^−^ phenotype [2]. The availability of quantitative information about CSCs dynamics allows to test pathways of their formation, resistance, and repopulation following irradiation using mathematical modeling approach.

The most used methods to model the growth of tumor cells, including CSCs, and their response to therapeutic effects, including radiation, are ordinary differential equations (ODEs) (for example, [3 – 5]), partial differential equations (for example, [6]), as well as agent-based methods, including cellular automata models (for example, [7]). ODE models describe the growth dynamics of a tumor cell population in terms of volume, number of cells, or concentration of cells or molecules. In some cases, variables representing cells or tissues other than cancerous ones are included, but their meaning is similar.

The mathematical model of a tumor (glioblastoma) was proposed in [4], incorporating two subpopulations of cancer cells undergoing transitions from CSCs to non-stem cells, but not *vice versa*, i.e. not undergoing dedifferentiation. The authors showed the fundamental possibility of tumor repopulation after therapeutic exposure (in that case, immunotherapy) just due to the surviving CSC subpopulation. It should be noted that the authors of [4] did not verify their results with any experimental data.

The population of tumor cells is modeled in [8] as consisting of two types of cells, stem and non-stem, each of the subpopulations divides, dies (under radiation exposure) and transits into another type with its own rates. The transition of non-stem cells to senescence is not considered. In addition, both types of transition, from non-stem cells to stem cells and *vice versa*, in the model are not associated with asymmetric division. The article focuses on the search for optimal modes of radiation fractionation that ensure maximum death of CSCs, depending on the parameters of different cell subpopulations growth.

A different approach was applied in [6]. The tumor model is implemented in the form of a system of partial differential equations, taking into account not only temporal but also spatial dependencies of the number of cells. It takes into account the diffusion of oxygen from the surface into the tumor and changes in the status of cells (dividing, temporarily resting, necrotic) depending on the oxygenation level. In addition, that work does not consider discrete subpopulations of cells (stem and non-stem), but a single population in which each cell has a “stemness degree” from 0 to 1, which depends on the local level of oxygenation. Thus, the authors take into account the fact that stem cells are localized mainly in hypoxic regions of the tumor. The probability of cell death upon irradiation is modeled by the standard linear-quadratic dose function, where the alpha and beta parameters depend on the oxygen status of the cell, its stemness, and the rate of division. Investigating the effect of radiation on the tumor during radiotherapy, the authors observe an increase in the proportion of high-stemness cells over time due to the greater radioresistance of hypoxic and high-stemness cells compared with normoxic and low-stemness cells.

This paper presents a biological model of the growth of cancer cells *in vitro*, which includes replenishment of the CSC pool due to dedifferentiation from non-stem cancer cells. This process is sufficient for a quantitative description of experimental data [7] on the dynamics of irradiated MCF-7 breast cancer cells, including cancer stem and senescent cells.

## METHODS

To study the mechanistic basis of the increased fraction of the CSC phenotype observed in cancer cell populations after irradiation, a biological model was introduced, Fig.1. The model was based on the following assumptions:

1. The tumor consists of three subpopulations: differentiated cancer cells (CC), cancer stem cells (CSC) and nondividing cells being in the state of radiation-induced senescence (SC) [9].
2. CSCs can divide symmetrically and asymmetrically [10].
3. Differentiated cancer cells are able to dedifferentiate, i.e. to transit into CSCs [11].
4. The growth rate of the cell population decreases as its volume increases. Population dynamics is described by an exponential growth function adjusted for the maximum number (N_max_) of cells that can fit in a given volume due to space constraints and nutrient availability.

**Figure 1.**
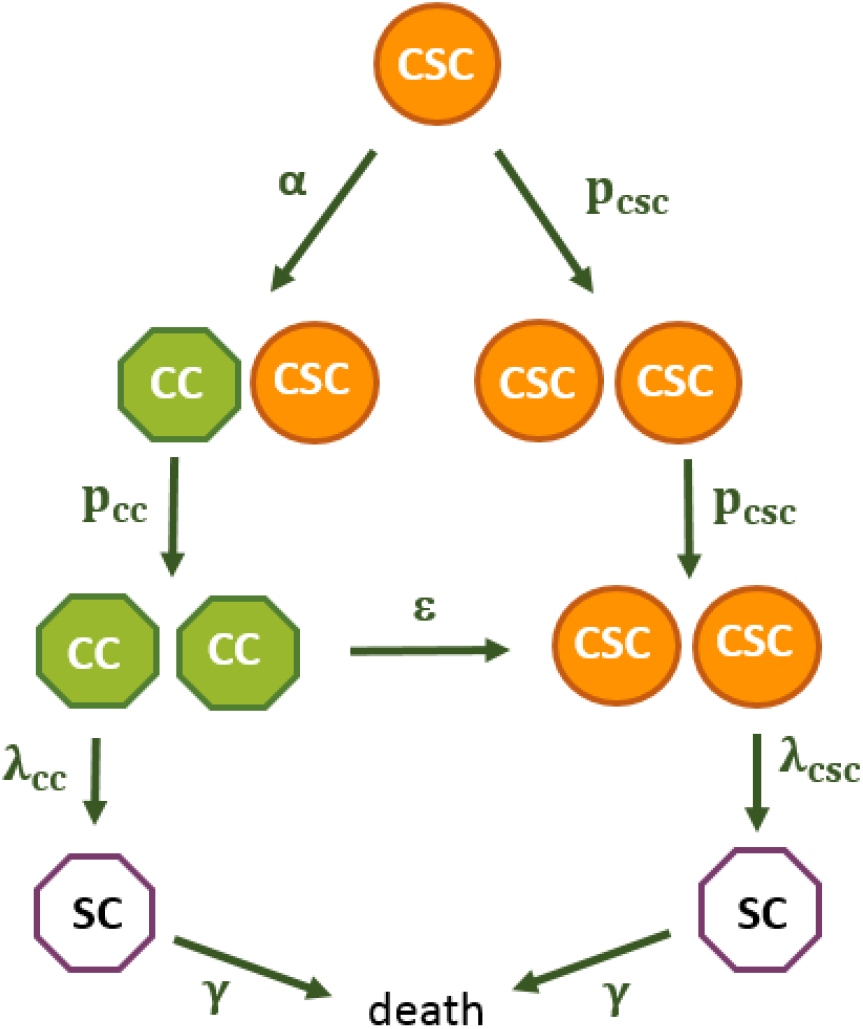
The scheme of the cancer cell proliferation model.

Both in control and under irradiation (single instantaneous irradiation), the dynamics of cancer cells is described by the following system of differential equations:

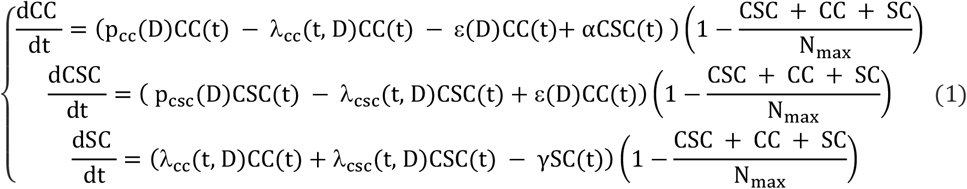

Here, CC is the number of differentiated cancer cells, CSC is the number of cancer stem cells, p_cc_(D) is the rate of proliferation of CCs, p_scs_(D) is the rate of symmetrical proliferation of CSCs, and *α* is the rate of asymmetric proliferation of CSCs. SC is the number of cells in the senescence state, *λ*_cc_(t, D) is the rate of transition of CCs to the senescence state, *λ*_csc_(t, D) is the rate of transition of CSCs to the senescence state, *ε*(D) is the rate of dedifferentiation of CCs. Non-dividing cells are eventually removed from the population at a rate of γ.

The rates of transition to the senescence state depend on the dose and time as follows:

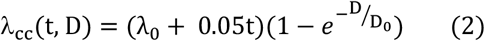

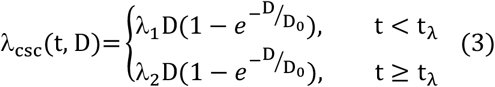

Here *λ*_0_, *λ*_1_, *λ*_2_, t_*λ*_, D_0_ are the parameters of the model. The rate of symmetrical proliferation of CSCs increases with dose as follows:

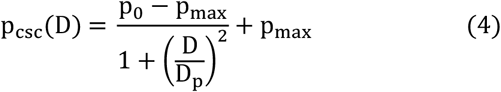

Here p_0_, р_max_, D_p_ are the parameters of the model. The rate of proliferation of differentiated cells decreases with dose increase as follows:

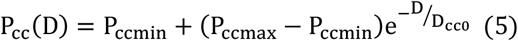

P_ccmin_, P_ccmax_, D_cc0_ are the parameters of the model. The rate of dedifferentiation depends on dose as follows:

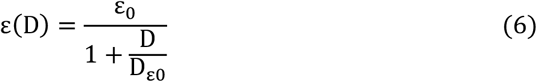

*ε*_0_, D_*ε*0_ are the parameters of the model. The model was implemented a) by numerically solving equations (1) using the MatLab package; b) using the Monte Carlo approach.

When studying the model by the Monte Carlo approach, a pool of cells is considered (the initial number of non-stem tumor cells is N_0cc_, stem tumor cells are N_0csc_). In a time step (Δt=0.005 days), each cell can undergo one of the processes shown in Fig.1, with probability Δt times the rate of the corresponding process. The model parameters were determined by the criterion of best agreement with the data obtained *in vitro* on MCF-7 cells [7].

## RESULTS

The simulation results in comparison with the experimental data [7] are shown in Fig. 2, the parameter values are shown in Table 1.

**Table 1.** The values of the model parameters obtained by the criterion of best agreement with experimental data *in vitro* [7], breast cancer cells MCF-7.

| Parameter | Value |
| --- | --- |
| $p_{ccmax}, \text{day}^{-1}$ | 0.95 |
| $p_{ccmin}, \text{day}^{-1}$ | 0.4 |
| $D_{cc0}, \text{Gy}$ | 0.3 |
| $\lambda_1, \text{day}^{-1}\text{Gy}^{-1}$ | 0.001 |
| $\lambda_2, \text{day}^{-1}\text{Gy}^{-1}$ | 0.09 |
| $t_\lambda, \text{day}$ | 5 |
| $\lambda_0, \text{day}^{-1}$ | 0.13 |
| $\gamma, \text{day}^{-1}$ | 0.001 |
| $D_0$, Gy | 0.3 |
| $\varepsilon_0$, day<sup>-1</sup> | 0.002 |
| $D_{0\varepsilon}$, Gy | 3.4 |
| $\alpha$, day<sup>-1</sup> | 0.01 |
| $p_0$, day<sup>-1</sup> | 0.2 |
| $p_{\max}$, day<sup>-1</sup> | 1.03 |
| $D_p$, Gy | 4 |
| $N_{\max}$ | 8e6 |
| $N_{0\text{csc}}$ | 720 |
| $N_{0\text{cc}}$ | 299280 |

**Figure 2.**
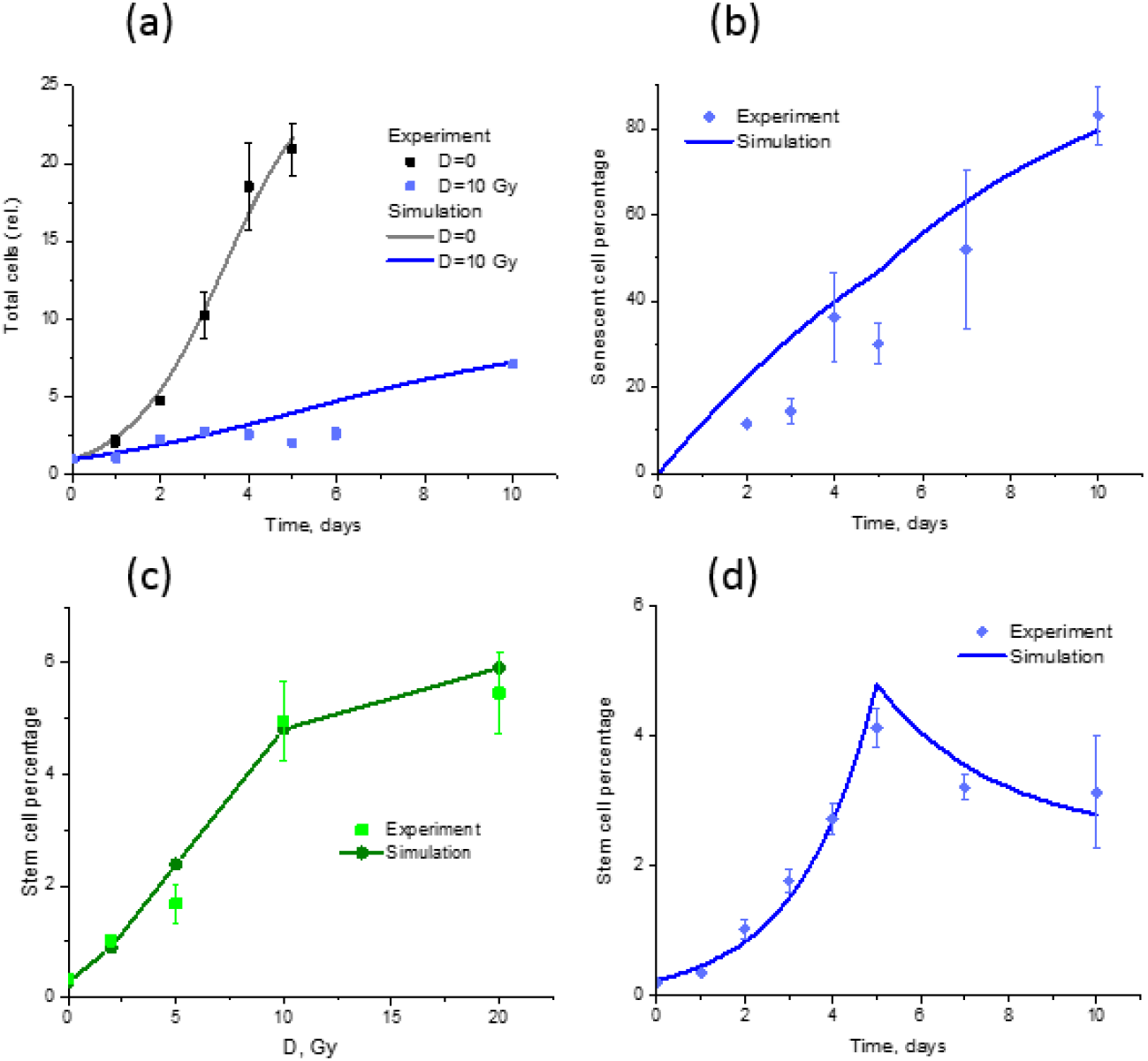
Description of *in vitro* experimental data [7], MCF-7 breast cancer cells, by the model. (a) Time dependence of the relative cell growth in control and after irradiation with a dose of 10 Gy. (b) Time dependence of the proportion of senescent cells in the population after irradiation with a dose of 10 Gy. (c) Dose dependence of the proportion of CSCs in the population on day 5 after irradiation. (d) Time dependence of the proportion of CSCs in the population after irradiation with a dose of 10 Gy. The values of the model parameters are shown in Table 1.

## DISCUSSION

The mechanisms ensuring maintaining of a pool of CSCs in a tumor and its repopulation after irradiation are widely discussed in literature [3, 4, 7]. Knowledge of such mechanisms is crucial for solving the problem of recurrence of malignant tumors after radiotherapy.

We analyzed the experimental data [7], Fig. 2, by the model comprising the dose-dependent dedifferentiation of non-stem tumor cells, as well as the dose- and time-dependent transition of tumor cells to a senescent state. We did not consider a subpopulation of pre-senescent cells and, accordingly, did not have an increase in the number of CSCs due to pre-senescent cells under irradiation [7]. The feature of our model is that the increase in the number of CSCs occurs only through symmetric division and dedifferentiation. This simplification turned out to be sufficient for explanation of data on both the dose and time dependence of cancer differentiated and stem cells after irradiation, as well as the time dependence of the senescence of MCF-7 cells.

A number of other works, for example [6, 8], raise the question in a more practical aspect, how to optimize the radiation therapy schedule in order to eliminate the maximum possible number of CSCs. However, various models do not look biologically verified enough, since their results, as a rule, were not compared with experimental data *in vivo* or *in vitro*. The biological ground of some models also raises questions. For example, in [4] no dedifferentiation of non-stem tumor cells to CSCs is considered, which is currently believed to play an important role in the repopulation of CSCs after irradiation [11, 12]. The model in [8] does not take into account the transition of non-stem cells to senescence, which may be involved in cell reprogramming [12].

A comparison with a number of existing studies shows that our simpler model is working as being demonstrated by the agreement of the calculated results with several experimental endpoints at once over a wide range of doses and times. In the future, a deeper incorporation of biological mechanisms is needed. Study of the role of the senescent state in the CSC repopulation may be important in further investigations.

## REFERENCES

1. Zheng D., Preuss K., Milano M.T. et al. Mathematical modeling in radiotherapy for cancer: a comprehensive narrative review. Radiat. Oncol. 2025, 20:49.

2. Al-Hajj M., Wicha M.S., Benito-Hernandez A. et al. Prospective identification of tumorigenic breast cancer cells. Proc Natl Acad Sci USA (2003), 100:3983–3988. doi:10.1073/pnas.0530291100

3. Weeks S.L., Barker B., Bober S. et al. A Multi-Compartment Mathematical Model of Cancer Stem Cell Driven Tumor Growth Dynamics. Bull. Math. Biol. 2014, 76: 1762–1782.

4. Abernathy K. and Burke J. Modeling the Treatment of Glioblastoma Multiforme and Cancer Stem Cells with Ordinary Differential Equations. Comput. Math. Methods Med. 2016, 1239861.

5. Swanson E.R., Kose E., Zollinger E.A., Elliott S.L. Mathematical Modeling of Tumor and Cancer Stem Cells Treated with CAR-T Therapy and Inhibition of TGF-β. Bull. Math. Biol. 2022, 84:58

6. Celora G.L., Burne H.M., Kevrekidis P.G. Spatio-temporal modelling of phenotypic heterogeneity in tumour tissues and its impact on radiotherapy treatment. J. Theor. Biol. 2023, 556:111248

7. Gao X., Sishc B.J., Nelson C.B. et al. Radiation-Induced Reprogramming of Pre-Senescent Mammary Epithelial Cells Enriches Putative CD44+/CD24−/low Stem Cell Phenotype. Front. Oncol. 2016, 6:138.

8. Meaney C., Kohandel M., Novruzi A. Temporal optimization of radiation therapy to heterogeneous tumour populations and cancer stem cells. J. Math. Biol. 2022, 85:51

9. Campisi J. Senescent cells, tumor suppression, and organismal aging: good citizens, bad neighbors. Cell 2005, 120: 513–22.

10. Peitzsch, C., Kurth, I., Ebert, N. et al. Cancer stem cells in radiation response: current views and future perspectives in radiation oncology. Int. J. Radiat. Biol. 2019, 95, 900–911.

11. Vlashi, E. and Pajonk, F. Cancer stem cells, cancer cell plasticity and radiation therapy. Semin. Cancer Biol. 2015, 31: 28–35.

12. Paul R., Dorsey J.F., Fan Y. Cell plasticity, senescence, and quiescence in cancer stem cells: Biological and therapeutic implications. Pharmacol. Ther. 2022, 231:107985.

